# How to maintain a high virulence: evolution of a killer in hosts of various susceptibilities

**DOI:** 10.1101/674994

**Authors:** Aurélien Chateigner, Yannis Moreau, Davy Jiolle, Cindy Pontlevé, Carole Labrousse, Annie Bézier, Elisabeth Herniou

**Affiliations:** BioForA, INRA, ONF, 45075, Orleans, France; Cirad. UMR ASTRE, Campus International de Baillarguet, 34398 Montpellier, France; Insect-Virus Interactions Group, Department of Genomes and Genetics, Institut Pasteur, CNRS; IRBI, UMR 7261, CNRS - Universite de Tours, 37200 Tours, France

## Abstract

Pathogens should evolve to avirulence. However, while baculoviruses can be transmitted through direct contact, their main route of infection goes through the death and liquefaction of their caterpillar hosts and highly virulent strains still seem to be advantaged through infection cycles. Furthermore, one of them, *Autographa californica* multiple nucleopolyhedrovirus, is so generalist that it can infect more than 100 different hosts.

To understand and characterize the evolutionary potential of this virus and how it is maintained while killing some of its hosts in less than a week, we performed an experimental evolution starting from an almost natural isolate of AcMNPV, known for its generalist infection capacity. We made it evolve on 4 hosts of different susceptibilities for 10 cycles and followed hosts survival each day. We finally evaluated whether the generalist capacity was maintained after evolving on one specific host species and tested an epidemiological model through simulations to understand how.

Finally, on very highly susceptible hosts, transmission-virulence trade-offs seem to disappear and the virus can maximize transmission and virulence. When less adapted to its host, the pathogen’s virulence has not been modified along cycles but the yield was increased, apparently through an increased transmission probability and an increased latent period between exposition and infection.

## Introduction

Survival of highly virulent pathogens is an old question brought back to the front scene since the 2013-2016 West-African Ebola epidemics (Sofonea et al. 2017). Understanding how highly lethal viruses can be maintained is also a leading question for agriculture and the development of biological pest killers. Virulence evolution theory predicts that pathogens should evolve to become avirulent as mild infections allow transmission over longer periods than fast host-killing lines (Méthot 2012). However, virulence can be adaptive and correlated to transmission and intra-host competition (Alizon and Michalakis 2015). For instance, it has recently been shown that Ebola infection through dead bodies was responsible for the transmission of the most virulent strains (Sofonea et al. 2017). However, this is not the most common transmission route for this virus, which is generally transmitted by direct exchange of fluids (Bausch et al. 2007). For baculoviruses, which infect pest caterpillars, transmission mainly occurs via direct ingestion of viral particles, on the form of occlusion bodies (OBs), which are released after the death and liquefaction of the previous host (Slack and Arif 2006). An individual baculovirus particle, such as the natural isolate of *Autographa californica* multiple nucleopolyhedrovirus (AcMNPV) carries many genetically diverse genomes, harboring different single nucleotide variations (SNVs), insertions or deletions (INDELs) (Chateigner et al. 2015; Gilbert et al. 2014, 2016). It is thus a population of individual virus genotypes. By analogy with Ebola, baculoviruses may maintain highly virulent lines through infection by contamination from dead bodies, therefore the virulence of mild populations adapting to a specific environment should increase (Sofonea et al. 2017).

According to the current transmission-virulence trade-off hypothesis and the Ebola case, a virus line transmitting through host death, perfectly adapted to its environment should evolve to reach an optimum of high virulence and high transmission rate. However, mechanically, if a host is killed rapidly, it means that it has less time to develop, limiting the carrying capacity of the host. Serial passage experiments in different hosts were shown to lead to virulence attenuation (Pavan, Boucias, and Pendland 1981). Ebert and Weisser (Ebert and Weisser 1997) showed it is not optimal for a pathogen to kill the host before its growth slows, even when the parasite is rapidly multiplying. For pathogens saturating their host cells, like baculoviruses, killing rapidly means that less resources will be available to produce OBs. Baculoviruses are known to delay their host molting by controlling the insect molting hormone (O’Reilly and Miller 1989, 1990), and thus delaying the host growth decreased speed during molting and allowing the virus to produce more. Thus, when reaching the optimum highly virulent state, a baculovirus line should also reach a trade-off optimal state of low yield. In contrast, when the pathogen is poorly adapted to the environment, more constraints can be met, like a lower susceptibility of the host requiring a higher number of particle entries to start an infection, or a lower reproduction rate inside the cells. The virus line can then adapt to the host and follow the previous scenario of high virulence, or stabilize around a low virulence and the particle production would be high, compensating the low virulence end ensuring the survival of the population. However, the question remains on the absolute levels of yield, whether a highly virulent line can produce a number of particles equivalent to low virulence strains.

AcMNPV is a generalist species, which is infectious to more than 100 different lepidopteran hosts(Cory 2003; Goulson 2003). However not all hosts are equally susceptible, so AcMNPV populations are subject to transmission-virulence trade-offs (Anderson and May 1982), which vary with the host species it may encounter in the wild. To understand the links between host susceptibility, virulence and yield, we formalized baculovirus infection process by adapting Sophonea epidemiological models and quantifying adaptive processes in hosts of varying susceptibility by experimental evolution. We expected our experimental design to cover the previously described cases of infection and to unravel the evolution of obligate killers.

## Materials and methods

### Virus and bioassays

We previously described our generation 0 virus population (Chateigner et al. 2015), obtained by in vivo amplification of an archival sample. AcMNPV-WP10 was extracted from a one-cycle infection of a large number of highly susceptible hosts (*Trichoplusia ni*), minimizing the selection pressure on viral genomes.

We had to choose the experimental evolution starting infection dose in order to maintain the infection for 10 generations and limit bias. In nature, the only infections that will be maintained on a long period have to start with a dosage at which viruses do not kill their hosts too fast, not having time to produce enough viruses for the next generation of infection and that do not lead to a non-killing infection, and thus a dead-end for the virus. We started bioassays to assess the proper dosage for our experiment and in our lab conditions, by testing the response of four caterpillar host species (*Trichoplusia ni, Spodoptera exigua, Manduca sexta* and *Agrotis ipsilon*) to our baculovirus AcMNPV-WP10. The four caterpillar host species were chosen because they represent a large range of susceptibility to the virus.

### The droplet method

The method used to infect the caterpillars is the droplet method (Li and Bonning 2007): a one night starved third instar caterpillar was isolated in the well of a plate and fed during 10 minutes with a 0.5 µL droplet containing the appropriate dose of virus, 4% (v/v) blue food coloring agent and 20% sucrose. Once the caterpillar was fed, it was transferred in an individual box with nutrient medium. Boxes were cleaned and the medium was changed every day until the caterpillar died or pupated. The dead caterpillars were transferred in a 1.5 mL tube, all the caterpillars from a same replicate where pooled together.

### Bioassays

We evaluated AcMNPV-WP10 original fitness in bioassays on the four different host species and with seven different doses: 50, 500, 2,500, 5,000, 7,500, 10,000 and 500,000 occlusion bodies (OBs) in the droplet. Three batches of 20 caterpillars were infected for each dose as biological replicates. The OBs yield has been counted on Thoma cells (in three replicates for 7 different concentrations when possible) after filtration on cheesecloth, two rounds of centrifugation (10 min at 7000 rpm) with SDS 0.1%, two other rounds with distilled water and then re-suspension in water.

After the experimental evolution described in the next paragraph, we estimated once again the fitness in bioassays on the four host species with the same method but only three doses, 50, 5,000 and 500,000.

### Yield, LT50, LD50 and survival

Bioassays allowed us to compute the components of the parasite fitness that we defined as yield and virulence, with proxies being the survival of the host, the lethal dose to kill 50% of the population (LD50), and the lethal time to kill 50% of the population (LT50). To compute yield, we averaged the multiplications of particles of each infection dose. For the LD50 and LT50, we used the dose.p function from the R package MASS with default parameters on a binomial model fitting the mean deaths of the replicates to respectively the log of concentrations or to the days after infection, with the glm function from the R package stats. The host survival was simply representing the number of hosts dying every day post infection.

### Evolution

#### Experimental evolution setup

To realize the experimental evolution, we infected 10 caterpillars of one species with a subsample of AcMNPV-WP10 G0 for 10 cycles: when the 10 caterpillars were dead, their bodies were pooled, the OBs were extracted (generation 1) and used to infect a new generation of caterpillars, and so on until the 10th generation. We put aside caterpillars that turn to pupae and did not count them as dead. We did not control the exact dose administered at each cycle, we chose to dilute the collected viruses by the yield of G0, specific for each host species. We did this work in 10 replicates for each hosts. We built 40 independent viral lines (figure 1).

**Figure 1:**
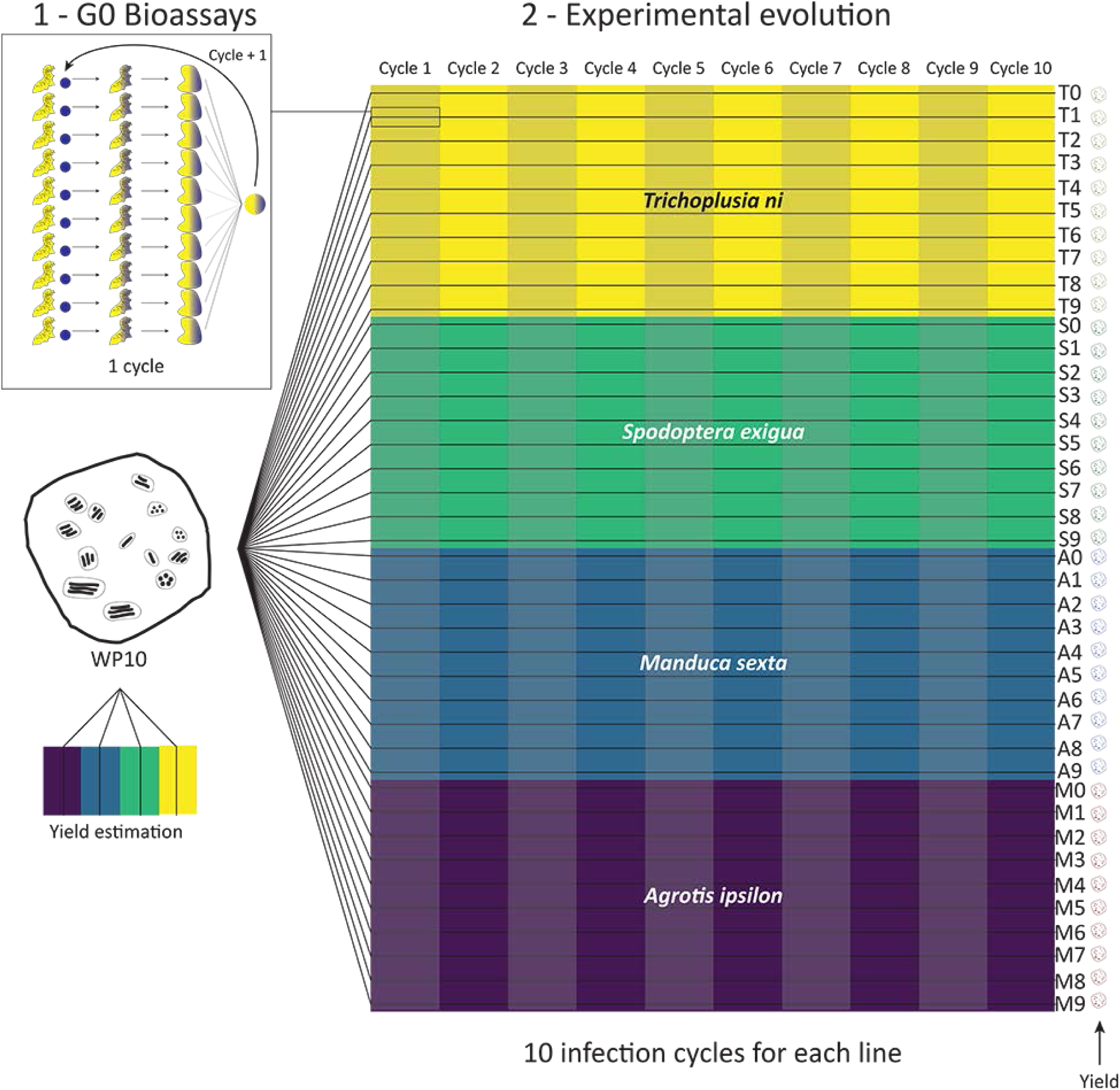
Design of the experimental evolution of the baculovirus AcMNPV-WP10 on four different host species. (1) Bioassays were realized by infecting 10 caterpillars by the generation 0 virus by the droplet method (Li and Bonning 2007). Once dead and liquefied, caterpillars bodies were pooled together and the generation 1 virus is extracted. It is then used to infect a new batch of 10 caterpillars. (2) From the original virus sample, 10 independent viral lines consisting of 10 infection cycles were created for each host species (*T. ni, S. exigua, A. ipsilon* and *M. sexta*) and for the mixed experiment, by the previously described protocol. The yield of the 40 lines (named T, S, A and M 1 to 10) is estimated in the end and compared to the original WP10 yield.

### Decomposition and forecasting of the lines

In order to properly represent, analyze and forecast the future path of each viral line, we treated them as classical time series of 10 cycles, where a cycle lasts for 23 days (22 days after the infection day). We used the R package forecast (Hyndman and Khandakar 2008) and specifically the mstl function, which decomposes time series. We used default parameters to compute trend, seasonality and remainder.

Our virus lines all come from the same original population. To account for this essential link, we treated them with hierarchical models, from the R package hts (Hyndman et al. 2011; Wickramasuriya, Athanasopoulos, and Hyndman 2018), and the function hts with default parameters. The top level of our model is the AcMNPV-WP10 starting population, the second level represents the caterpillar species, and the third level is the virus line one (figure 2).

**Figure 2:**
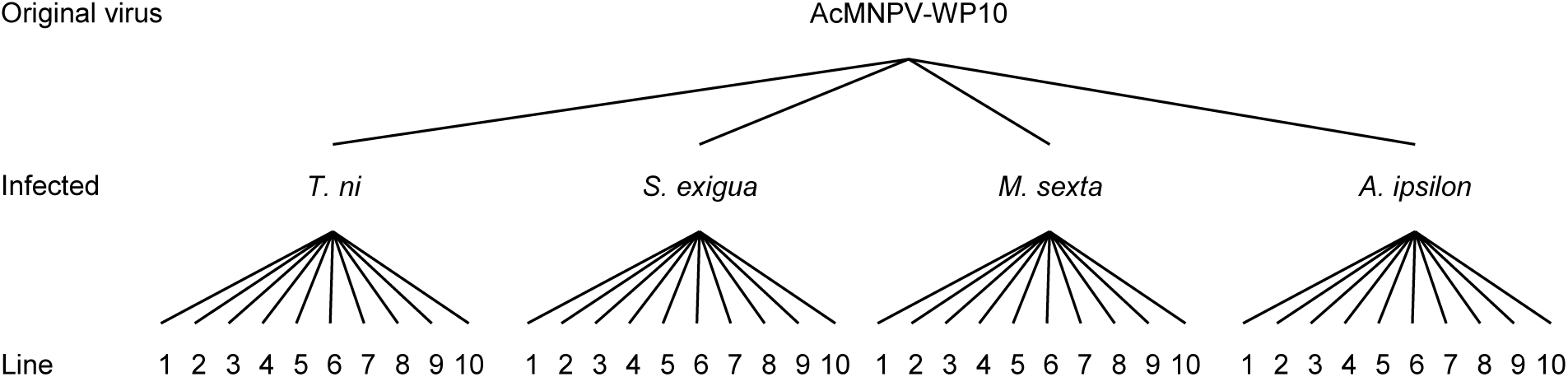
Hierarchical model used to forecast the future of virus lines.

As our counts of dead caterpillars were discrete with a high proportion of zeros, we cannot directly use quantitative models to forecast our data. Models specific to count data exist but did not perform well on our data, forecasting aberrant data. Thus, in a first step, we transformed our data by multiplying them by 100, and we added 1 to be able to constrain forecasts to an interval. This way, the minimum value of 0 is transformed to 1 and the maximum value of 10 is transformed to 1001. We constrained the data to a 0 - 1002 interval with the formula presented in equation 1 and de-transformed the forecasts with the formula presented in equation 2.

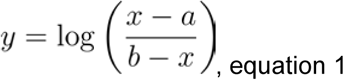

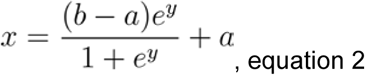

Where y is the transformed data, x is the data on the original scale, a and b were respectively the lower and upper boundary of the constrained space.

#### Epidemiological model

Our SEIRD model, presented in figure 3, is a classical extension of the Susceptible-Exposed-Infected-Recovered (SEIR, Keeling and Rohani 2011) model with a “death” (D) compartment except that like in the Ebola virus SEICRD model (Sofonea 2017), this last compartment contains an infected dead body class that allow for post-mortem transmission (Legrand 2007, Weitz & Duschoff 2015, Sofonea 2017). Furthermore, direct transmission remains possible but is marginal and due to accidental events like cannibalism, thus baculovirus transmission is mechanically different from spore transmission. It is an extreme case where post-mortem transmission is the main way of infection.

**Figure 3:**
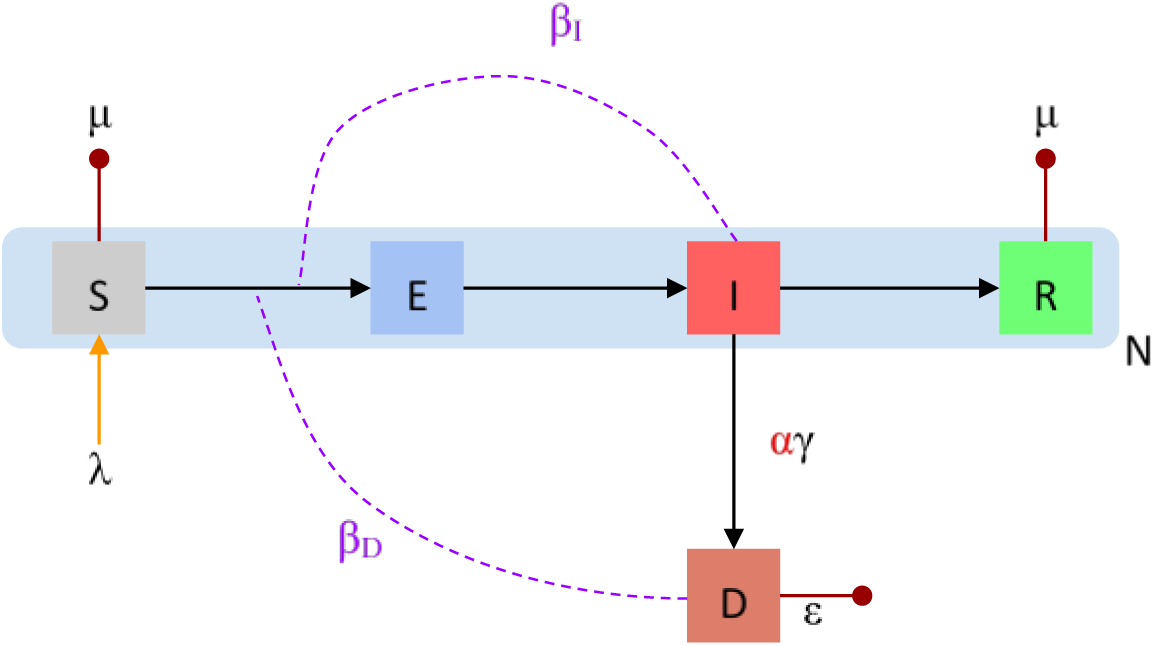
Epidemiology of baculoviruses. S, E, I, R and D were respectively the susceptible, exposed, infected, recovered and death compartments. λ is the host inflow, μ is the natural host death rate, ω is the infection rate, α is the virulence, γ is the arms race period and ε is the rate at which dead bodies loose infection capacity. βI and βD were adapted from the SEICRD model and were the rates at which infected caterpillars and dead bodies infect new individuals. Finally, N is the size of the population. In our experiment, βI is controlled and equal to 0.

The experiment starts by the infection of N caterpillars at t_0_ (N_0_ = N). No more host inflow is further allowed (λ_0_ = N, λ_>0_ = 0). The susceptible (S) proportion of the population always represents 100% of the population as we only keep in the experiment caterpillars infected by the droplet method. The natural host death rate μ is equal to 0 during the whole experiment (verified in a control experiment, no caterpillar died when fed by the same method with virus replaced by water in the droplet) before molting.

For the first infection cycle, all the susceptible caterpillars become exposed then infected at a rate ω = 1, as we control that each caterpillar actually ingests the virus, which dose was chosen to infect 100% of the caterpillars. This rate will vary during the next infection cycles as viruses evolve and manage to adapt to the environment. It represents the time needed by the baculovirus to penetrate the midgut epithelium of the caterpillar and usually represents 24 hours.

In the nature, infected caterpillars can directly infect other caterpillars by accidental events like cannibalism, at a rate β_I_. However, this rate is of low magnitude and is controlled to be null in our experiment as caterpillars were isolated. The infected caterpillars leave this compartment at a rate γ. This parameter referred to as the inverse of symptomatic infectious period by Sofonea *et al.* would be the equivalent to the arms race period between the host and the virus. Caterpillars finally die (D) from the virus at a rate αγ - where α is our measure of the virulence - or recover (R) at a rate (1 - α)γ. The dead bodies infect the next generation at a controlled rate of β_D_ = 1. They were infectious for a period ε^−1^. In our experiment, the infections were controlled and chained with only a few days between cycles in order to maintain infectivity of particles. However, we did not measure the loss of infectivity and we thus chose a secure infectious period of 120 days as we keep our viruses at −20°C (Jorio, Tran, and Kamen 2006).

Our model does not include any additional host heterogeneity as we chose the hosts to be as homogenous as possible.

Simplifying the basic reproduction number from the SEICRD model, we found at the disease-free equilibrium, the number of secondary infections caused by a single infected individual in a fully susceptible population (Diekmann 1990) is

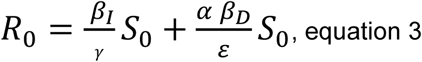

Where S_0_ is the total population size at the beginning of the experiment, β_I_ is the direct contact transmission, γ is the rate at which infected caterpillars leave the infected compartment, a is the virulence, β_D_ is the post-mortem transmission factor and ε is the inverse of post-mortem infectious period. As we isolated caterpillars during the whole experiment, we only considered the killer transmission (β_I_ = 0) and we can simplify equation 3 to

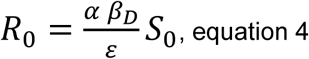

#### Model simulations and evaluation

We estimated model parameters for generation 0 and 10 by simulating the model for 23 days with a grid search approach: we arbitrarily defined possible values for each parameter and tested all the combinations, and then we computed the root mean square error (rmse) and the mean absolute error (mae) between the simulations and the bioassay for which we searched parameters. The simulation with the lowest rmse and mae was selected to represent the bioassay.

The grid is the following:

| Parameter | Possible values |
| --- | --- |
| Dead bodies contact rate | 50, 500, 2500, 5000, 500000 |
| Transmission probability | From 0 to 1, by 0.1 |
| Infectious period | 0.001, 0.01, 0.1, 1, 2, 4, 5, 7, 10 |
| Latent period | 0.00001, 0.0001, 0.001, 0.01, 0.1, 1, 2, 4, 10 |
| Arms race period | From 1 to 10 |

With $\beta_D$ = Dead bodies contact rate x Transmission probability, $\omega$ = 1 / Latent period, $\gamma$ = 1 / Arms race period and $\alpha$ = 1 / Infectious period,

## Results

### Characteristics of the original population

To have the best chances to go through 10 cycles of infection in the different host species, we carefully characterized the virulence of our original population for these specific environments. The first bioassays provided the resistance characteristics for the host species : *Trichoplusia ni* is the most susceptible species to WP10, with an LD50 of only 52 particles and LT50s of only 5.25, 4.92 and 1.85 days at respectively 50, 5000 and 500,000 particles in the infection droplet, and a yield 135 times higher than the infection dose (figure 4**A**); *Spodoptera exigua* is the second most susceptible, with the highest production of 273 times more particles than the infection dose, but also higher LD50 of 280 particles and LT50s of 14.85, 6.65 and 4.89 at the same doses. For the two remaining species, the order is less clear and depends on the trait studied. For yield and LD50, *Agrotis ipsilon* is more susceptible than *Manduca sexta*, with 3 times more particles produced versus 1.4, and 24,400 particles being sufficient to kill 50% of the population versus an imputed dose of 44 million, which is too high to be put in a 0.5 µL droplet. For LT50, the difference is not clear, as no *A. ipsilon* caterpillar died when infected by 50 and 5,000 particles, but the LT50 is of 7.5 days at 500,000 particles, while *M. sexta* caterpillars died from 5,000 particles, but the LT50 is of 19.62 days, and still as high as 11.42 when infected by 500,000 particles. With these results, we defined the *T. ni* and *S. exigua* as species susceptible to WP10 and *A. ipsilon* and *M. sexta* as resistant species. We thus chose to start the experimental evolution by infecting our caterpillars at 2 different doses, 2500 particles for the low resistance species and 500,000 particles for the high resistance species.

**Figure 4:**
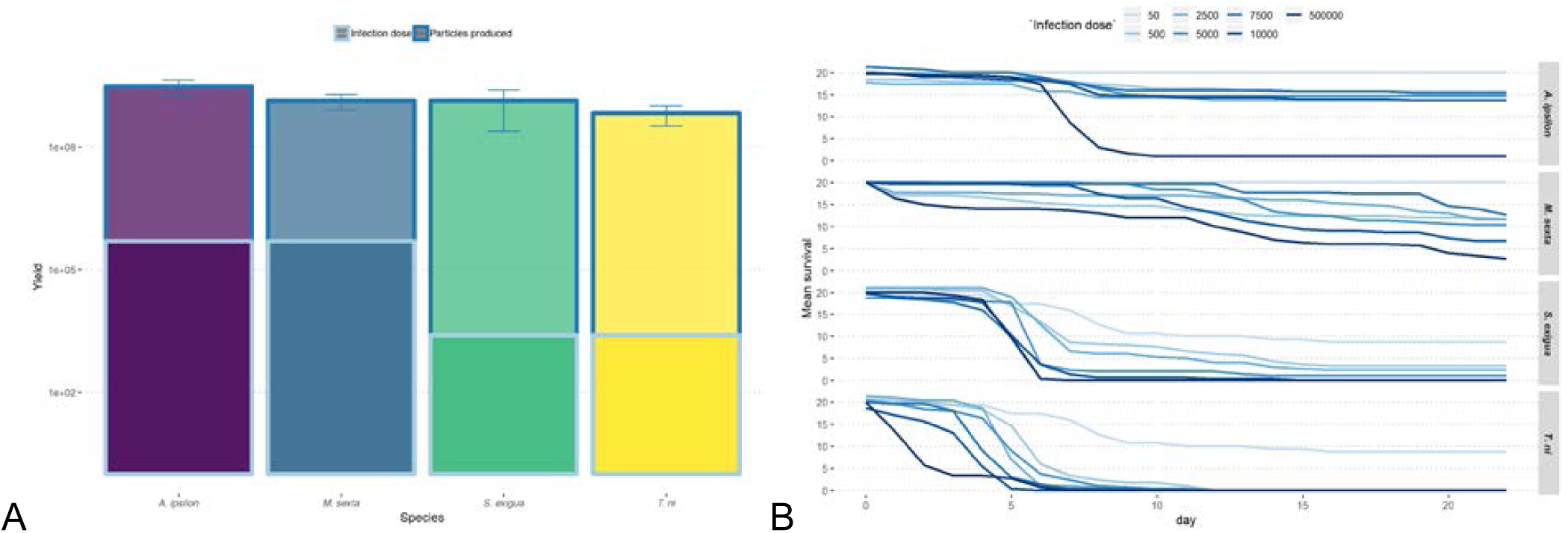
(A) *A. ipsilon, M. sexta, S. exigua* and *T. ni* caterpillar production of particles after infection by *AcMNPV*-WP10 G0 compared to the infection dose and (B) survival to the virus for 7 doses, along the 23 days of the experiment.

The mortality of these lines, presented in figure 4**B**, along the 23 days of the experiment showed different resistance patterns for the different host species. *T. ni* caterpillars were very susceptible to the dose of infection and consistently killed by the virus from low doses. *S. exigua* ones showed smaller dose-response, but were killed consistently by the virus. *M. sexta* caterpillars showed a stronger resistance to the virus and a high correlation between dose and response. Finally, *A. ipsilon* caterpillars were only killed efficiently when infected by a very high dose.

### Ten cycles of evolution

We treated the experimental evolution infection cycles as time series, in order to be able to describe the evolution of each line but also to resume it for each species. With a classical decomposition of the data in trend, seasonal and remainder, we were able to represent each line trend, to compute mean trends and variances per host species (figure 5).

**Figure 5:**
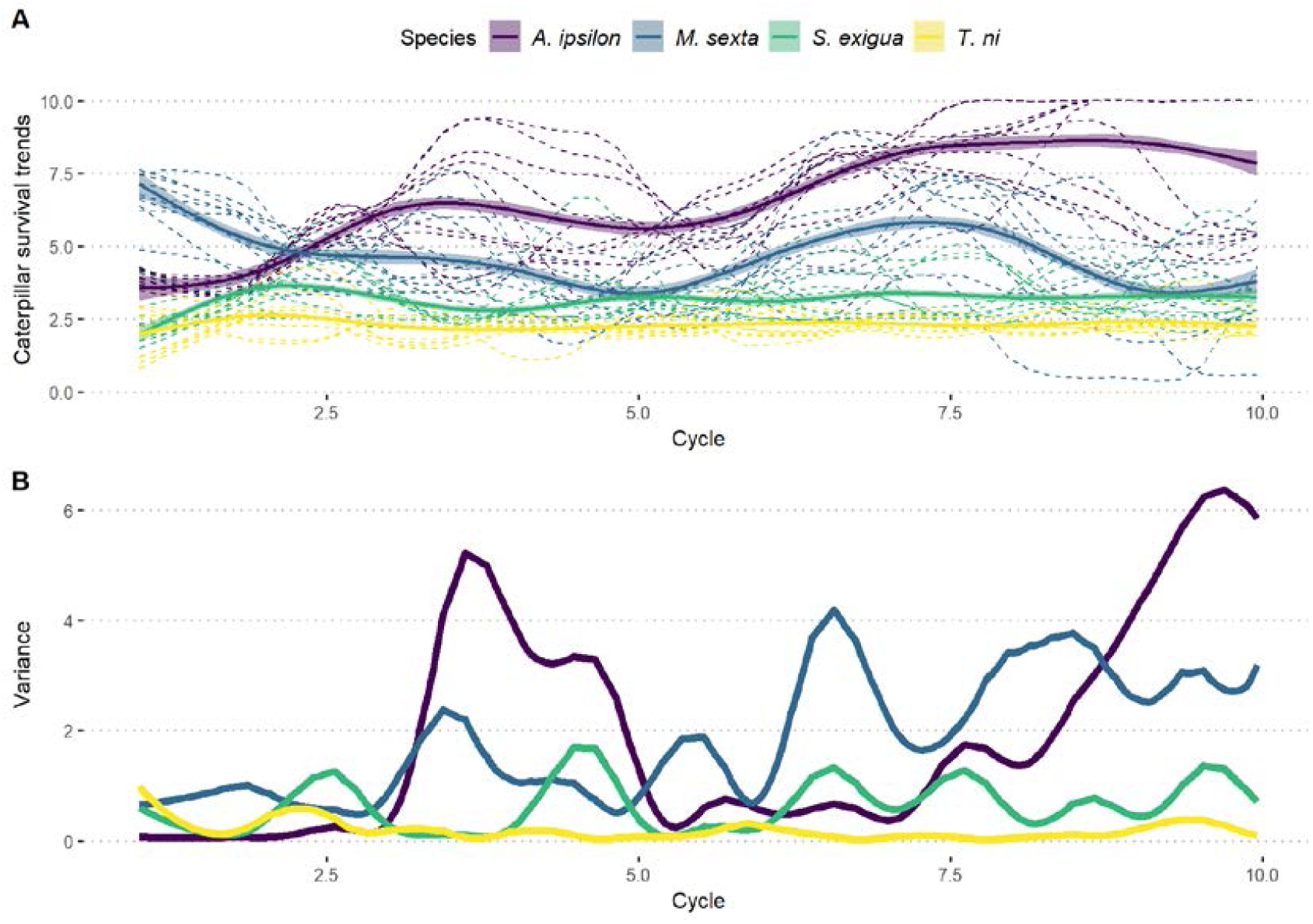
Trends and variance of caterpillar survival along the experimental evolution. (A) Plain lines represent mean trends for each species, with ranges representing 99% confidence intervals and dashed lines for individual lines. (B) Plain lines represent variance between viral lines evolving on same species along experimental evolution.

Viruses that evolved on *T. ni* maintained a very low caterpillar survival along cycles, and those that evolved on *S. exigua* maintained a slightly higher but stable caterpillar survival. When evolving on *M. sexta*, viruses were able to kill caterpillars with varying efficiency, as the mean trend was oscillating. However, on *A. ipsilon*, the global trend shows that viruses were not able to kill efficiently caterpillars along the cycles as survival increases (figure 5A). Moreover, there is more variation between lines that evolved on *A. ipsilon* than on *M. sexta, S. exigua* and *T. ni* (respectively 4.76, 3, 0.68 and 0.18 caterpillars, figure 5B). While this variation is stable and low on *T. ni* and *S. exigua*, it is increasing along the cycles for the two others. Only four lines that evolved on *A. ipsilon* were able to reach the 10th cycle, the six other lines had lost any virulence to the caterpillars between cycles 6 and 8 (figure S1).

**Figure S1:**
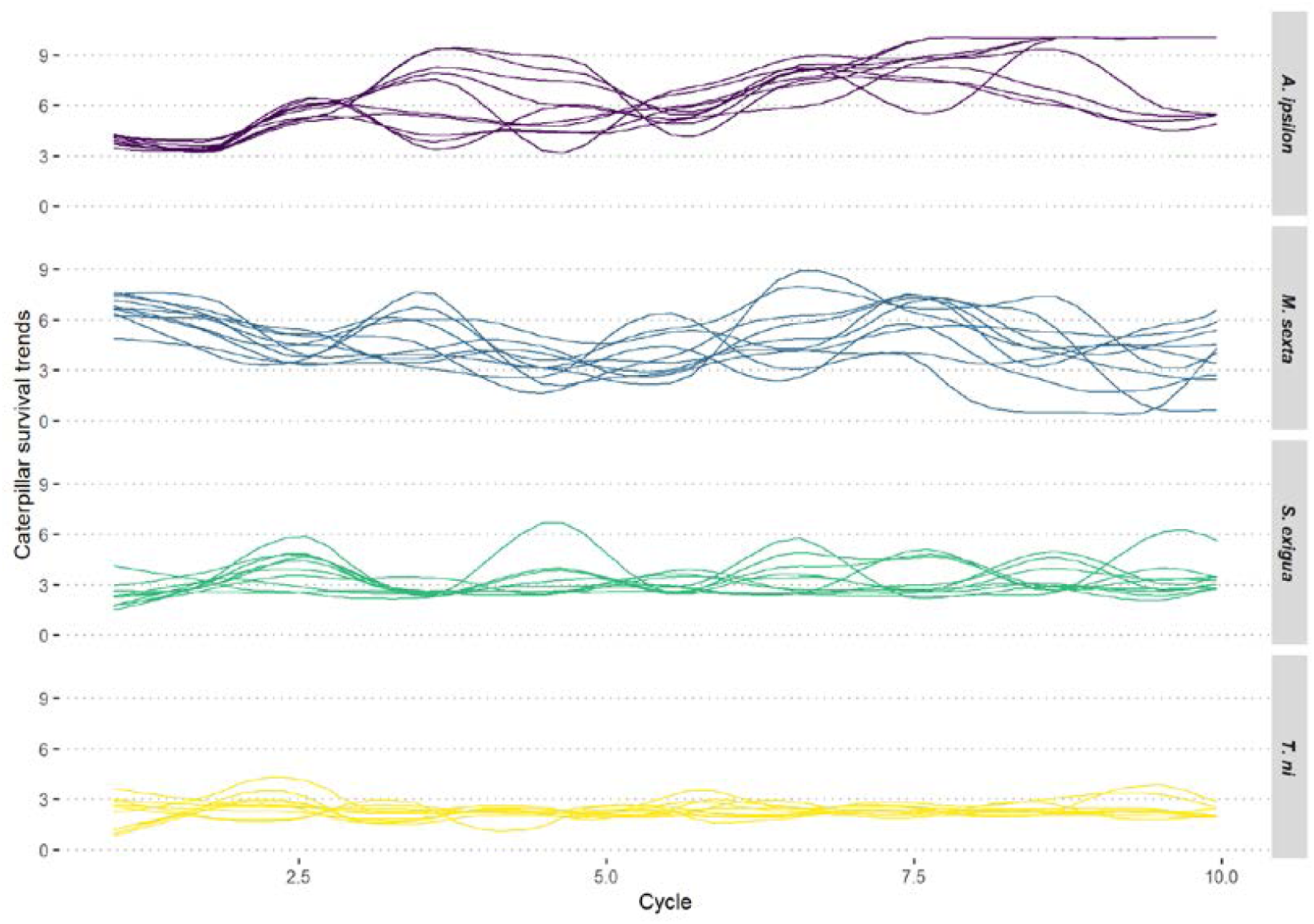
Individual caterpillar survival trends of the different lines during experimental evolution for the four host species.

### Evaluation of virulence after evolution

After the experimental evolution, we were able to appreciate how the different virus lines evolved in the unique host species and in the unstable environment (figure 6). While its yield increased, the host that was producing the highest number of particles per caterpillar, *S. exigua*, became the second producer behind *T. ni* lines. While yield slightly increased on *M. sexta*, the other resistant species (*A. ipsilon*) produced less particles after 10 generations of evolution than after the first cycle. After evolution we tested the virulence of *T. ni* and *S. exigua* lines to the host on which they evolved, but also to the other hosts (figure 7, figure S2). We were not able to test for the other 2 species as we were not able to harvest enough virus particles in last generations to run the experiment. However, differences appeared between the susceptible hosts lines within each host species, between host species and compared to the original population. It appeared that lines that evolved on *T. ni* were, on average and on this same host, slightly less virulent in the first days at 500,000 particles but more virulent at 5,000 and 50 particles. For *S. exigua* lines on this same hosts, the overall virulence was very similar to G0’s, even if they presented a less steep slope. However, lines that evolved on *T. ni* showed a virulence equivalent to the G0 on *S. exigua*; On the other hosts, the variance between lines was very high and it was thus difficult to rely on per species description. Lines that evolved on *T. ni* tended to be more virulent at low dose but less virulent at high dose on *M. sexta*; and they were less virulent on *A. ipsilon*. Lines that evolved on *S. exigua* were less virulent on *T. ni*, on *M. sexta* and on *A. ipsilon*, even if they showed capacity to kill few *M. sexta* caterpillars at 50 particles. The main difference in the infection of the more resistant caterpillar species, between lines that evolved on *T. ni* and lines that evolved on *S. exigua*, was that the former were always able to kill more caterpillars at the end of the experiment.

**Figure 6:**
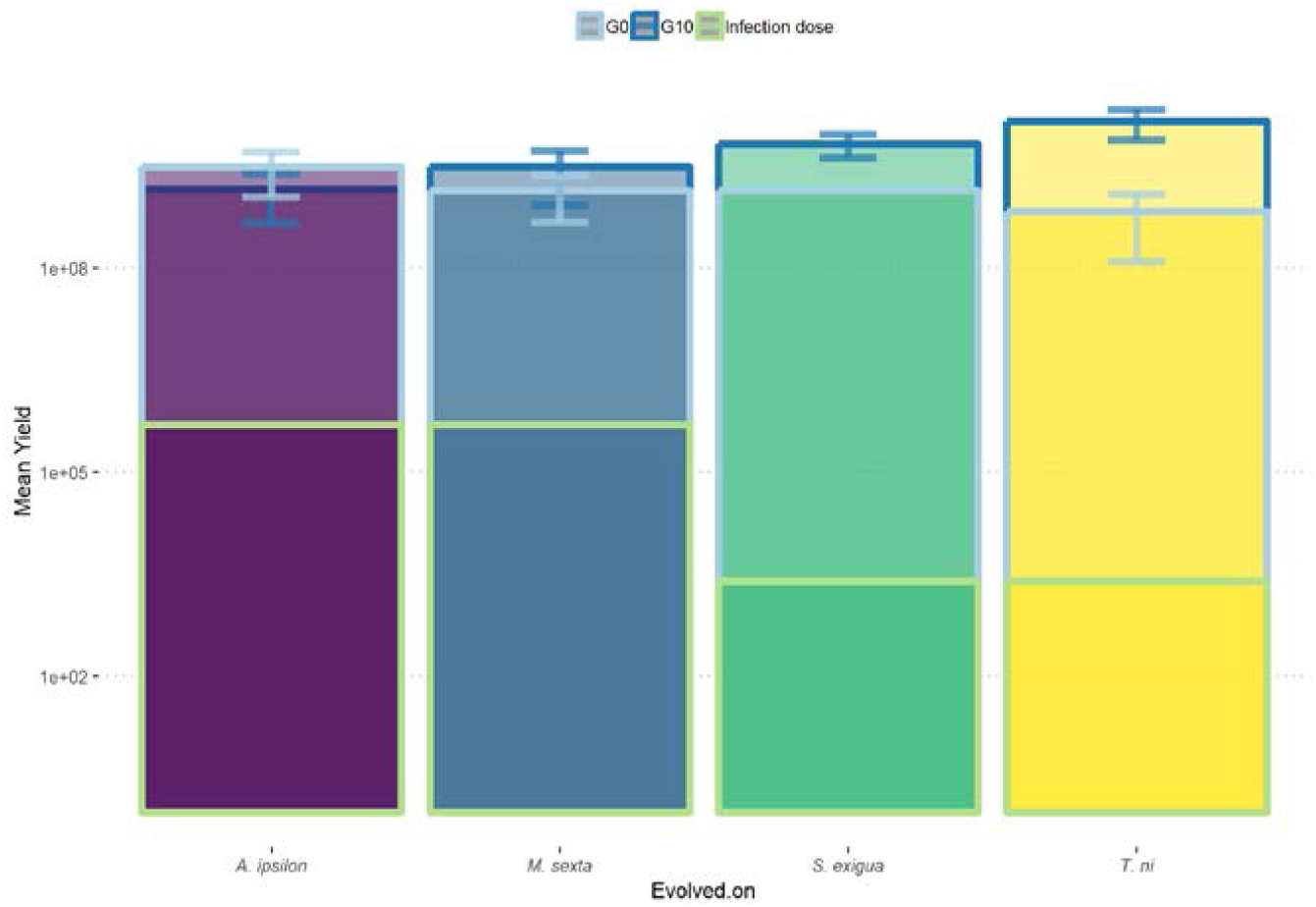
*A. ipsilon, M. sexta, S. exigua* and *T. ni* caterpillar production of particles after infection by *AcMNPV*-WP10 G0 and G10s compared to infection dose for G0.

**Figure 7:**
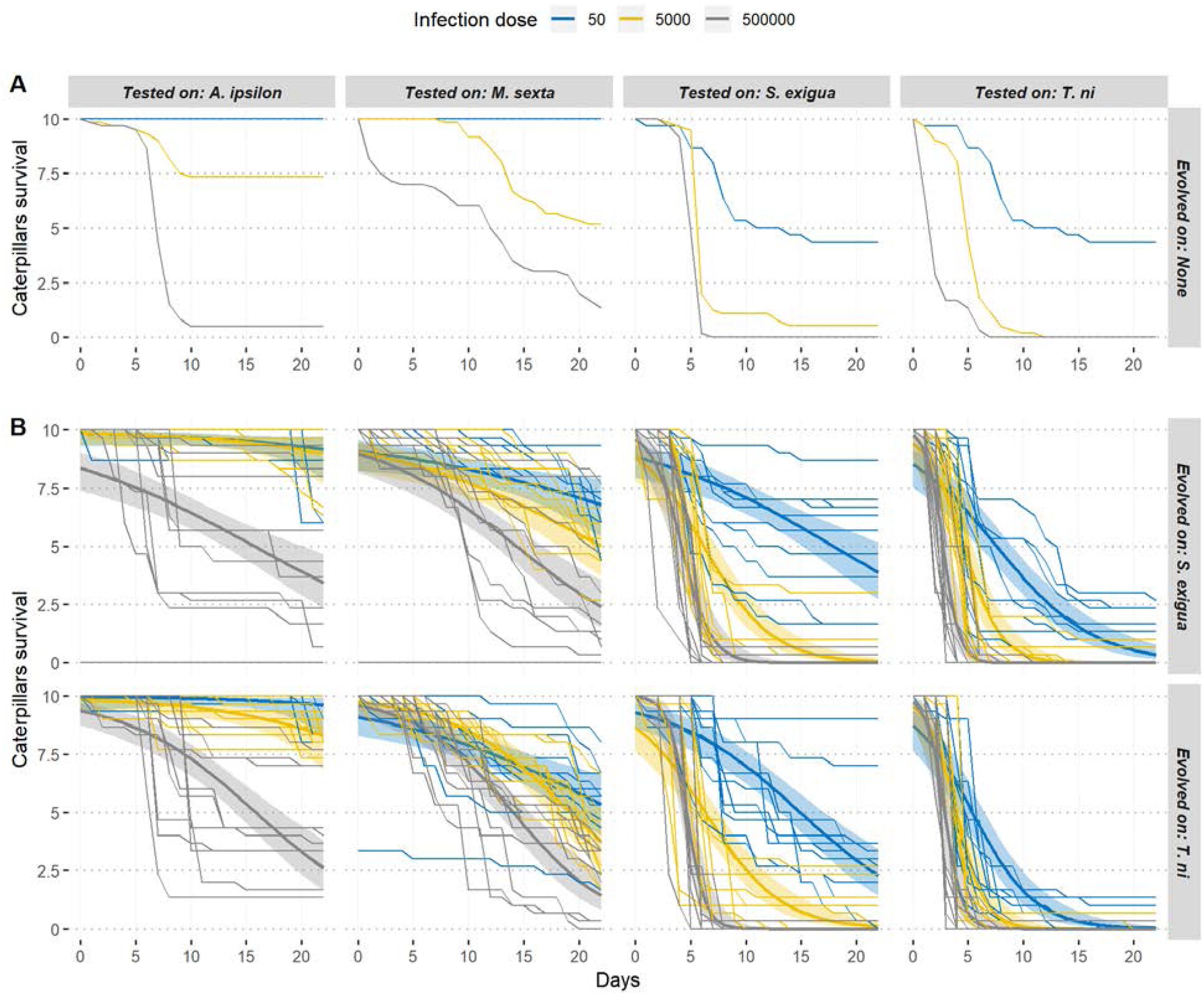
Morality of *A. ipsilon, M. sexta, S. exigua* and *T. ni* caterpillars from virus lines that did not evolve (G0, panel A), evolved on *S. exigua* or on *T. ni* caterpillars (panel B, respectively upper and lower part), at the 4 different doses used. The smooth lines represent the fitted binomial curve with standard deviation around it. The other lines represent the raw mortality due to the different lines.

Individually (figure 7), the most striking results was that while lines that evolved on *T. ni* were consistently and rapidly killing caterpillars of this same host, from the lowest dose, lines that evolved on *S. exigua* did not show this capacity: no line was able to kill all the *S. exigua* caterpillars at the lowest dose and there is even one line that was still not able to kill 75% of the caterpillars at 5000 particles. Another point of interest was that lines that evolved on *T. ni* were able at low dose to kill *S. exigua* caterpillars more efficiently than ones that evolved on *S. exigua*. When dose increases, the results of these lines became very similar to lines that evolved on *S. exigua*.

**Figure S2:**
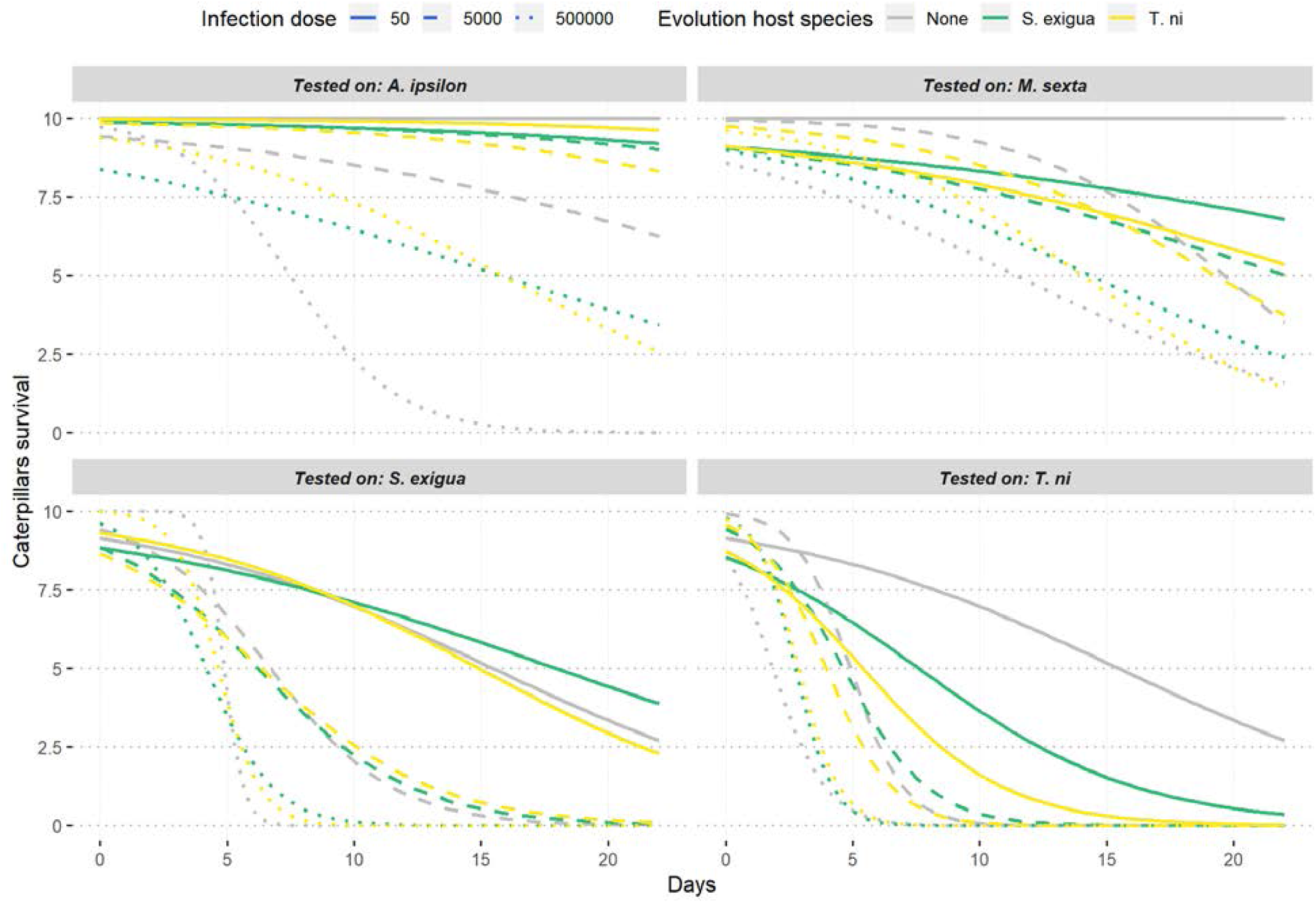
Mean mortality of *A. ipsilon, M. sexta, S. exigua* and *T. ni* caterpillars from virus lines that did not evolve (G0, in grey), evolved on *S. exigua* (in green) or on *T. ni* caterpillars (in yellow), at the 3 different infection doses.

### Evolution of the fitness

We compared the estimated fitness of the evolved lines to the original one (figure S3). Basically, fitness of the original population shows a steeper reaction norm than evolved lines at the 3 tested values. On *T. ni* and *S. exigua* caterpillars, evolved lines showed the exact same median R0. On *M. sexta*, results vary according to the tested dose, while it is clear on *A. ipsilon* that lines evolved on *T. ni* have a lower fitness than ones evolved on *S. exigua*. However, evolved lines showed the exact same fitness for the highest dose, always equal or lower than the original line. However, the evolution of the parameters estimated provided a more detailed information on the processes by which evolution happens in the different compartments of the epidemiological model (figure 8). When tested on *T. ni* caterpillars, the evolved lines showed results consistent with the original population. On *S. exigua* caterpillars, arms race period and transmission probability both increased. On *M. sexta*, it is the latent period and the transmission probability that increased. Finally on *A. ipsilon*, all the parameters drastically increased.

**Figure 8:**
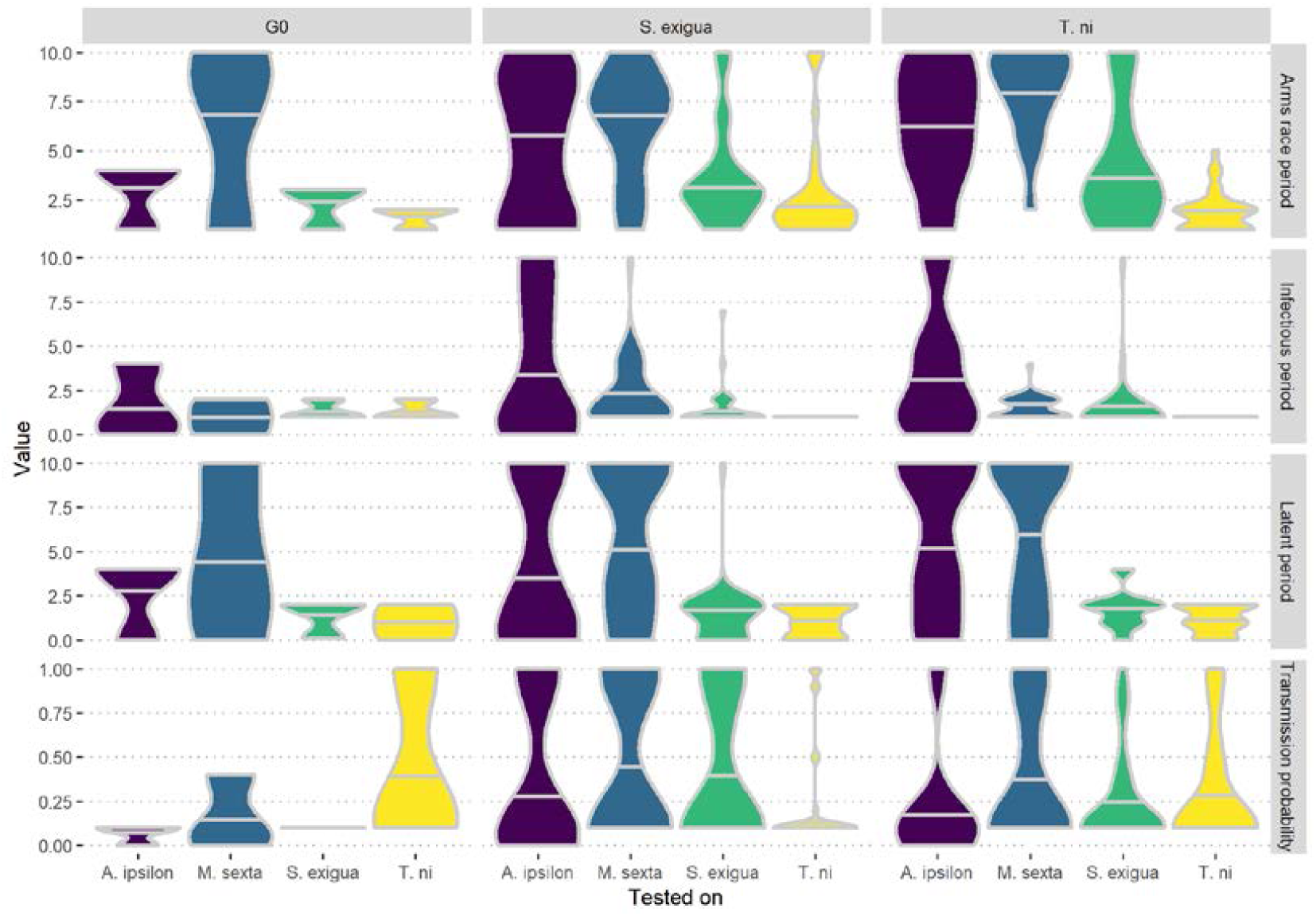
Distribution of the 4 different parameters (arms race period, infectious period, latent period and transmission probability) of the epidemiological model estimated in the simulations, for the ancestral (left column) and evolved virus lines (on *S. exigua*, middle column; on *T. ni*, right column), tested on the 4 different host species.

**Figure S3:**
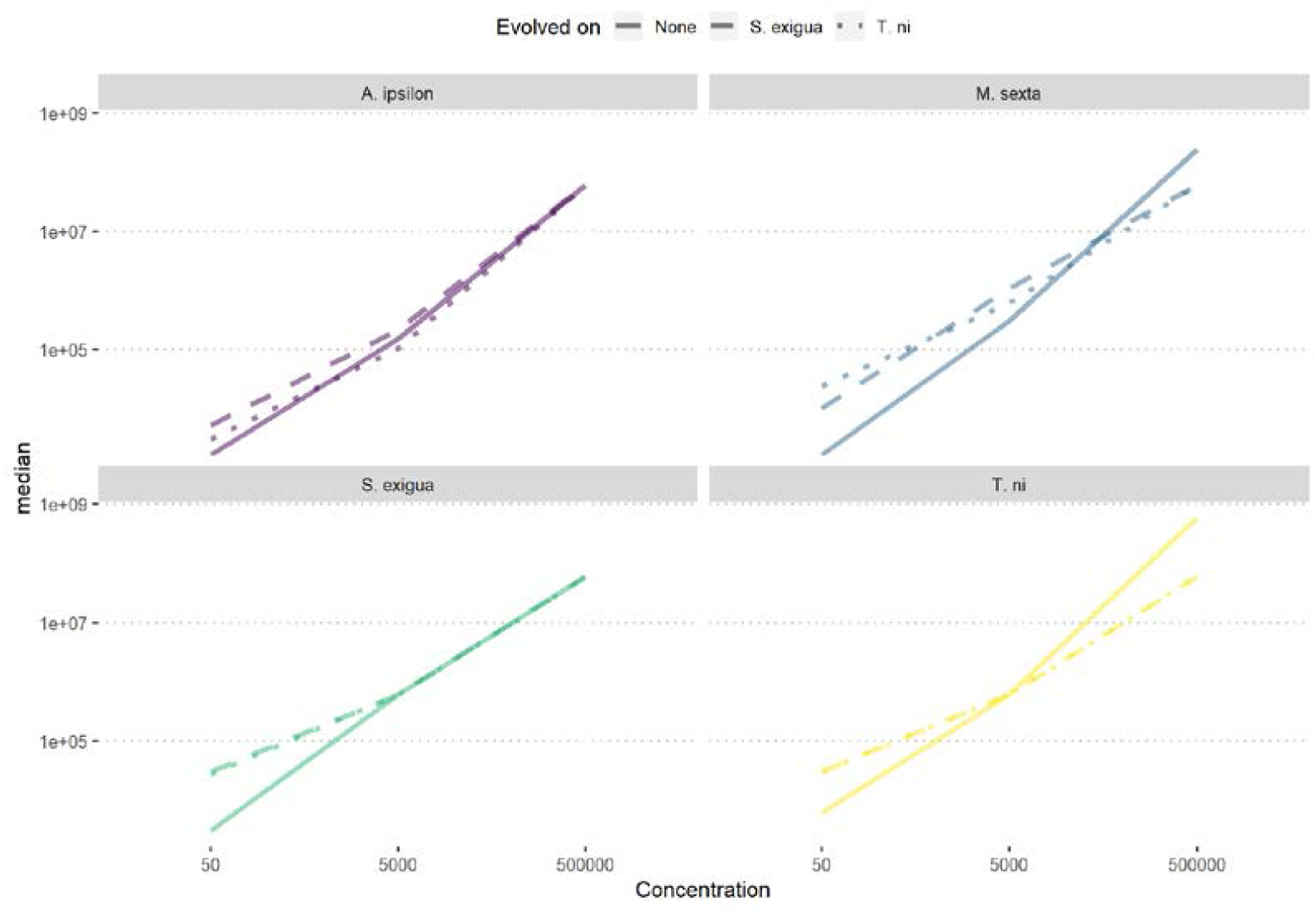
G0 (“None”, solid lines) and median evolved lines (*S. exigua*, dashed lines and *T. ni*, dotted lines) R0 estimated on *A. ipsilon, M. sexta, S. exigua* and *T. ni* caterpillars (respectively in purple, blue, green and yellow), for the 3 infecting doses of 50, 5,000 and 500,000 particles.

## Discussion

AcMNPV-WP10 is a generalist virus with the potential to infect a large number of host species, with a wide range of virulences for these different hosts. It is mainly transmitted through dead bodies infection, but a direct transfer between individuals remains possible. These characteristics make it an interesting model to study the evolution of viruses that can be transmitted through dead bodies like Ebola, but which are not limited to obligate killing transmission. It is also interesting directly for its use as a biopesticide to understand how the virus can maintain in caterpillars populations and remain virulent.

Before the experiment detailed in this manuscript, our virus population has been sampled from nature and has suffered on cycle of replication in *T. ni* caterpillars in order to multiply the number of virus particles available (Li and Bonning 2007; Chateigner et al. 2015). The main concern with this amplification was that any better performance of the virus on this host may be due to this amplification and would thus only be artifactual. However, the baculovirus original population was isolated from one caterpillar of another species (*Autographa californica*) and the virus did not show equivalent virulence on any other species on which we made it evolve. This experiment thus confirms that of our 4 host species, *T. ni* is the most susceptible to the virus. In the virus community, virus species are named after the host species from which it has been isolated. However, this method is criticized because a generalist virus can be found on a host that is not the natural reservoir and thus generating inaccurate names. We were not able to obtain *Autographa californica* caterpillars to compare our virus virulence on this host because of European import laws and thus exhort our american colleagues to compare the virus performance on both hosts.

During evolution, we were able to represent how strong and stable the infection is on *T. ni* caterpillars, as the mean of the trend is equal to 2.30 caterpillars surviving and very little variation appears during the cycles (0.181). On *S. exigua*, which is the second most susceptible species, the mean is, while slightly less, also low (3.19 caterpillars surviving) and stable (var = 0.684). This work resolves a previous interrogation as before evolution, we were not able to determine which of the *M. sexta* and *A. ipsilon* species was the most resistant host to the virus, as the results were not consistent depending on the character considered. On *M. sexta*, the survival mean trend was lower and more stable than on *A. ipsilon* (respective means of 4.65 and 6.81; respective variances of 3 and 4.76). Evolution showed that the virus is not able to maintain for 10 generations on *A. ipsilon* species consistently, while it survived on *M. sexta*. For the four virus lines that survived on *A. ipsilon*, their mean trend was equal to 6.22 and their variance was equal to 2.59. The ones that did not survived had a higher mean of 7.2 and a very high variance of 5.83. This large variance, expressing the instability of the infection from one infection cycle to the other may be the main reason of the collapse of these virus lines. If a threshold exists determining if a virus line can adapt to a host or not, repeating this experiment with more replicates could improve our estimations.

We thus think that *A. ipsilon* cannot be considered as a species susceptible to AcMNPV, as the virus cannot survive for a long time, even if the virus seems to be adapted to the host (R_0_ > 1, (Gandon et al. 2013)). After evolution, the estimated parameters (figure 8) of the epidemiological model showed a global increase, which did not happen in the hosts for which the host was adapted. We thus postulate that this global increase reflects the poor adaptation.

*M. sexta* caterpillars are growing faster than the other species of this study and to a higher body mass. The arms race in these caterpillars was thus more intensive in this species, as confirmed by the epidemiological model (figure 8). It may be an explanation for the slope of the decreasing phase of the seasonality, which was less steep than for the other species (figure S4). Furthermore, while killing caterpillars late when they reach a large size would produce a high number of virus circulating particles, we did not observe it during the experiment, caterpillars that died were always of smaller size than asymptomatic ones of equivalent age and days post infection. Thus, only relatively – for this species as they were killing more slowly than *T. ni* or *S. exigua* lines – fast-killing viruses were able to spread in these virus lines.

After the experimental evolution, virus that evolved on one host were expected to have changed their virulence and yield on the host on which they evolved, as selection pressure would select better adapted viruses. These characters should also be modified for the other hosts, as a side effect of the adaptation to the host: they were expected to specialize. However, if selective pressure were weak, only drift and intra-host competition would make the generalist potential to be modified. The latter would favor the viruses killing faster or yielding more circulating particles, as they would have more chances to be transmitted to the next generation. We can also expect that there is a maximum killing speed, mechanistically limited, and in that case, it would be possible for intra-host competition to overcome the trade-off between virulence and transmission and select the virus that optimizes host cells resources to increase yield, while being the fastest.

On *T. ni, S. exigua* and *M. sexta* caterpillars, after 10 cycles of evolution, we saw that yield increased compared to the original virus population, a lot more on *T. ni*, more on *S. exigua* and a little more on *M. sexta*. This increase follows the susceptibility order of host species. On the opposite, on *A. ipsilon*, the yield was lower after evolution. Lines did not adapt to this host and were collapsing.

We were not able to perform the last bioassays with the 40 lines for multiple reasons, like the quantity of viruses that we could extract that was too low for certain lines in the last generation, problems for rearing the caterpillars or synchronize them, and also because these experiments were very time consuming and projects are not eternal. We thus choose to focus on the lines that had the best chances to give results that we could trust, *T. ni* and *S. exigua* lines. It showed that less selection pressure was applied in *T. ni* lines than in *S. exigua* lines, as the former were able not only to kill its evolution host efficiently, but also to kill the other hosts tested more efficiently than the latter, especially at low dose. While globally most lines specialized for these environments, some lines were not able to adapt and lost infection capacity.

It is interesting to note that even the high infection doses used to infect *A. ipsilon* and *M. sexta* could not compensate for the weak adaptation of the virus to these hosts. We also did not control exactly the infection dose between the cycles and only diluted the extracted virus solutions by a factor specific to each species, which was the multiplication factor of viruses after the first infection experiments. This way, we were mimicking the natural stochastic spread of the virus and natural variations in infection dose.

In this article, we showed that if a virus evolves in an environment for which it is highly adapted, like in our experiment on *T. ni*, it can reach the highest virulence but also increase its yield. The classical transmission-virulence trade-off disappears with selection pressure and thus intra-host competition being the only selecting strength, it will improve the virus characteristics without any cost. We expect that the only limit to the pace of improvement is the mutation rate, which would require genomic studies beyond the scope of this article. Drift can still stochastically modify these characteristics, but with a weak impact. With a higher selection pressure like in *S. exigua*, intra-host improvements were reduced and specialization has stronger effects. When a population struggles to survive to an environment, but is able to survive, as we saw on *M. sexta*, the virulence did not change, but the yield slightly increased. It thus seems that the trade-off factor was decreased along the cycles. However, this conclusion would require keeping a more detailed track of virulence and yielding at each cycle. Finally, when survival of the population is at risk, because of maladapted virus, virulence seems to be the limiting factor. Indeed, for baculoviruses, killing the host implies that the virus has spread to the whole host before exploding the cells and liquefying the caterpillar. Thus, the capacity to infect all the cells and thus the virulence is the limit and this is what happened to lines evolving on *A. ipsilon*.

In conclusion, the transmission-virulence trade-off could be a transient phenomenon on the adaptation scale. From no adaptation to complete adaptation, virulence first has to reach a high enough value before the trade-off starts and finally disappears with complete adaptation.

**Figure S4:**
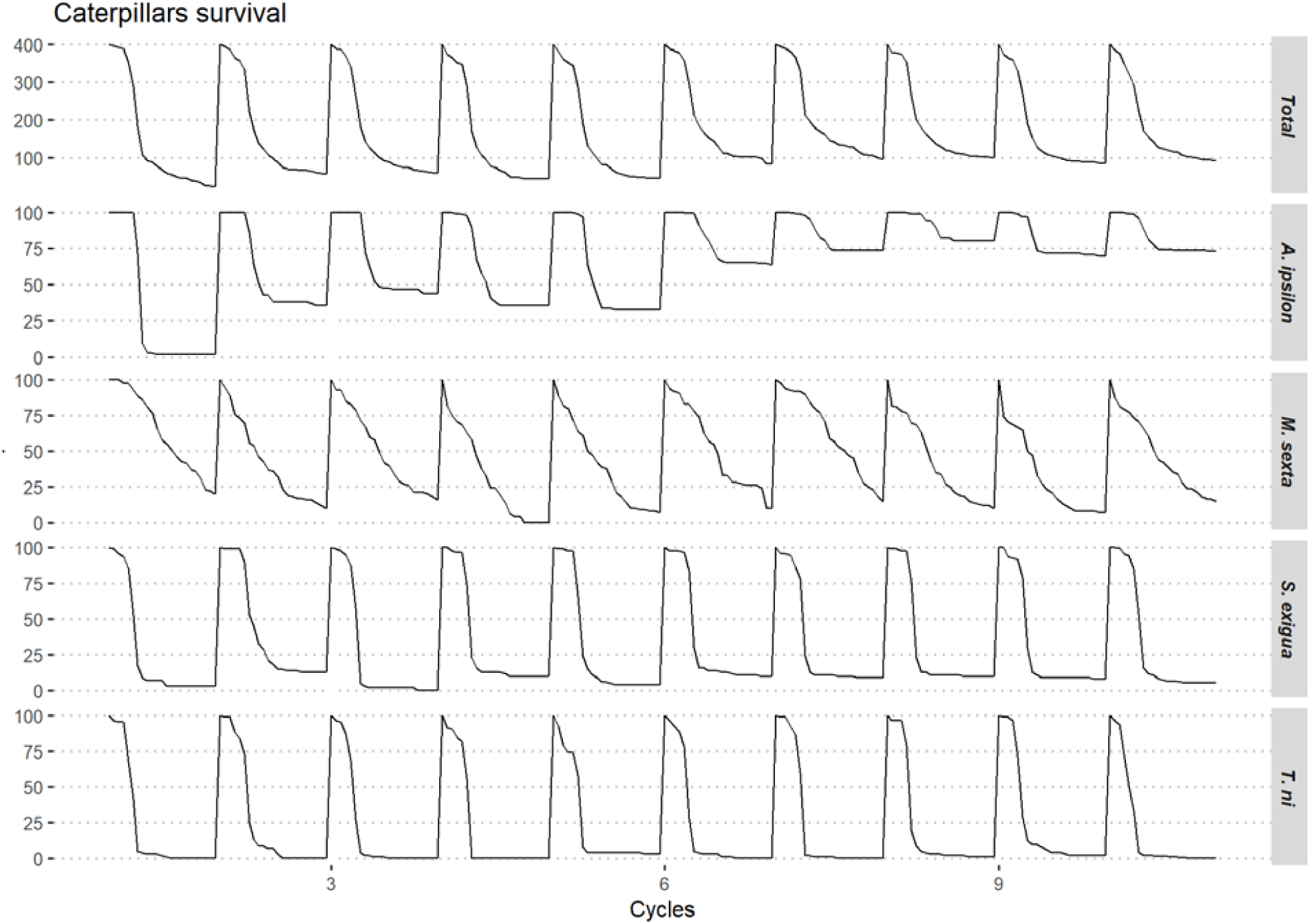
total (first row) and per species (the next 4 rows) aggregated survival for 10 cycles.

## Acknowledgements

The authors want to thank Samuel Alizon and Mathieu Tiret respectively for the discussions on epidemiological models and time series analyses.

## Funding

This work has been supported by the European Research Council (starting grant: Genovir 206205) and the CEA-Genoscope (project AP08/09#19).

## Statement of authorship

AC and EAH designed the experiment, discussed the results and wrote this manuscript. AC ran the *in silico* experiment. All authors ran the *in vivo* experiment.

